# A single-nucleus multimodal framework reveals epigenomic priming of chemoresistant states in ovarian cancer

**DOI:** 10.64898/2025.12.04.692102

**Authors:** Yuna Landais, Juliette Bertorello, Marta Puerto, Agathe Kahn, Baptiste Simon, Amélie Roehrig, Adeline Durand, Marceau Quatredeniers, Constance Lamy, Enora Laas, Nicolas Pouget, Simon Dumas, Jean-Guillaume Féron, Virginie Fourchotte, Claire Bonneau, Vincent Cockenpot, Marianne Delville, Magali Richard, Maud Kamal, Fabrice Lecuru, Christophe Le Tourneau, Céline Vallot

## Abstract

Non-genetic intratumor heterogeneity (ITH) drives therapeutic failure in cancer, yet its clinical monitoring remains challenging. We develop a multimodal single-nucleus framework that simultaneously profiles transcriptional and histone modification landscapes from frozen biopsies. Applied to longitudinal samples from 16 patients with high-grade serous ovarian cancer (HGSOC), this approach reveals reproducible tumor evolution under chemotherapy: proliferative and interferon-responsive states are lost, while those associated with TNFα- and epithelial–mesenchymal transition (EMT) expand. Chromatin profiling shows that these chemoresistant programs are epigenetically primed through H3K4me1 marks before treatment and nominates transcription-factor drivers, including ZBTB7A. Non-genetic baseline tumor composition predicts survival, and fibroblast remodeling parallels malignant adaptation. These findings establish a clinically scalable strategy for mapping functional ITH and identify epigenomic priming as a determinant of therapeutic failure.

**Graphical abstract:** 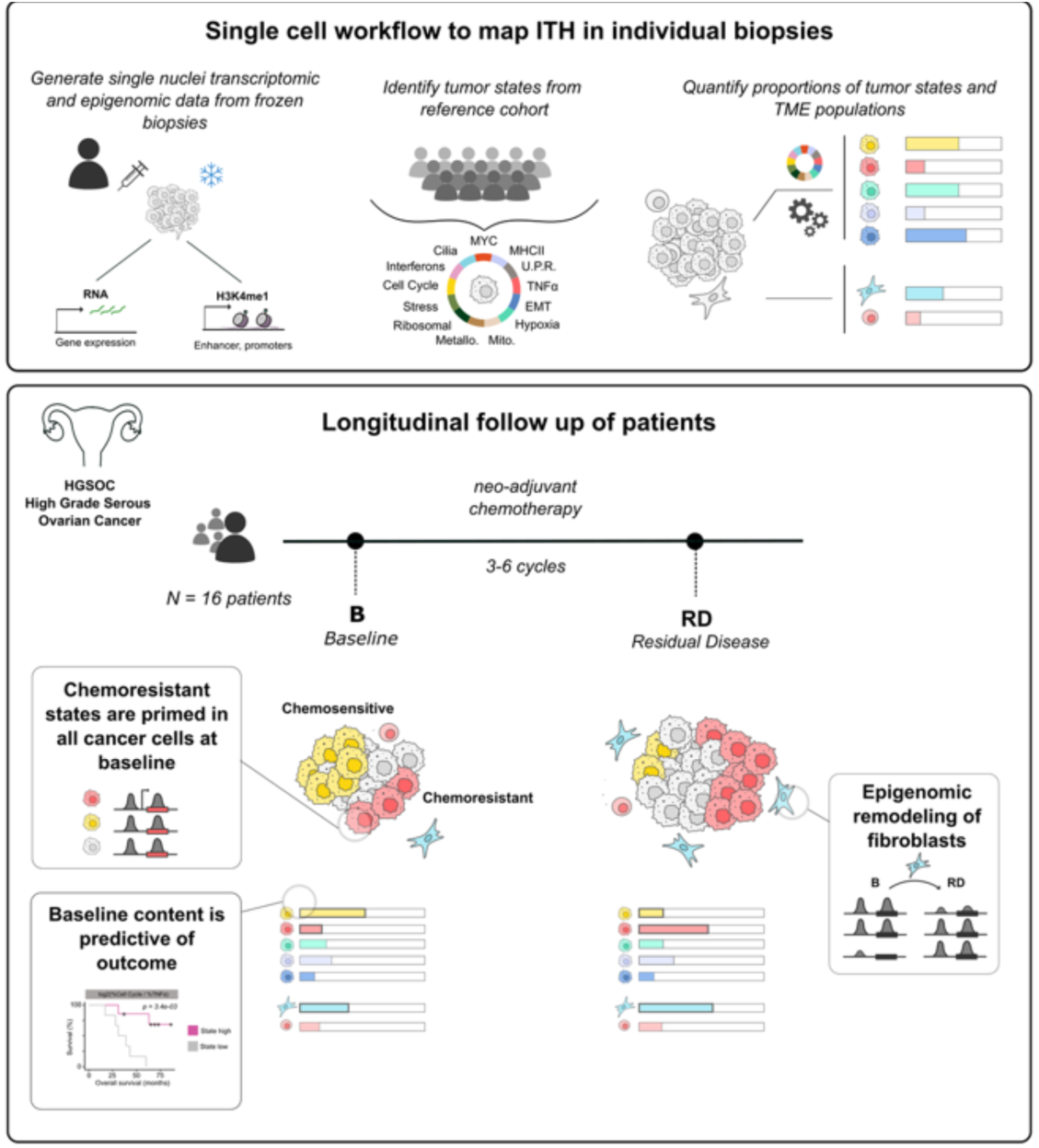

## Introduction

High-grade serous ovarian cancer (HGSOC) remains one of the most lethal malignancies, with most patients relapsing after platinum-based chemotherapy and a five-year survival below 40% ^1^. This poor outcome is due largely to intratumor heterogeneity (ITH), defined as the coexistence of genetically, transcriptionally, and functionally distinct tumor and stromal populations within the same tumor microenvironment (TME) ^2^. Such a complex mosaic of phenotypes drives adaptive resistance and immune evasion, preventing durable therapeutic responses. Understanding how this heterogeneity arises and evolves in patients is central to advancing precision oncology.

Clonal genetic evolution shapes long-term tumor progression^3^; increasing evidence shows that non-genetic variation, encoded in chromatin and transcriptional plasticity, can drive rapid adaptation of tumor cells to external stimuli, such as therapeutic stress ^4,5^. Epigenomic heterogeneity could provide a reservoir of metastable cell states that can be selected by therapy or microenvironmental cues ^6^. Pre-existing epigenomic configurations have been shown to fuel tumorigenesis in mouse models in the pancreas ^7–9^ and mammary gland ^10^, where a unique chromatin state primes pre-neoplastic cells and might be selected for throughout malignant evolution. Similar observations of epigenomic priming have been reported in studies of metastatic progression in a mouse model of lung cancer ^11^. Nonetheless, direct evidence for such an epigenetic priming in human tumors remains limited, largely because longitudinal molecular profiling at sufficient resolution has not been feasible in clinical practice.

Single-cell genomics has begun to map the regulatory programs defining malignant identity across human cancers ^12,13^. Most single-cell transcriptomics studies rely on freshly dissociated tissue, which limits their applicability for longitudinal or retrospective clinical cohorts. In contrast, single-cell epigenomic methods, such as single-nucleus (sn) ATAC-seq, can be performed on frozen patient material but primarily capture chromatin accessibility ^14–16^, providing limited resolution for the histone-based regulatory code that defines cell states ^17^. Single-cell histone modification profiling in cancer has remained challenging to use for patient material ^6,18^ and is mainly limited to individual use cases ^19^ or to model systems with higher input material ^20–23^. The ability to capture both transcriptional and epigenomic information from routine frozen biopsies remains a key barrier to translating single-cell discoveries into the clinic.

Here, we establish a clinically compatible multimodal single-nucleus approach that jointly profiles transcriptional and histone modification (H3K4me1) landscapes from milligram-scale frozen biopsies. This workflow overcomes a key barrier to patient-level epigenomic analysis by enabling high-resolution quantification of functional ITH directly from routinely collected material. By integrating large public scRNA-seq datasets with longitudinal samples from 16 patients with HGSOC, we generate a patient-resolved atlas of tumor and stromal evolution under chemotherapy. Our results reveal that tumor composition evolves reproducibly during chemotherapy: cycling, hypoxic, and interferon-signaling cells are lost, while TNFα-signaling populations expand. The baseline balance of these cell states correlates with overall survival, with the presence of TNFα-dominant tumors correlating to poorer outcomes. In sum, this resource provides a framework for dissecting chromatin-encoded plasticity in human tumors and can be readily extended to other malignancies where frozen biopsies are available.

## Results

### Integrated single-nucleus profiling from frozen biopsies enables multimodal tumor analysis

We performed multimodal single-nucleus profiling on paired baseline and post-chemotherapy residual disease (RD) biopsies from 16 patients with high-grade serous ovarian cancer (HGSOC) enrolled in the SCANDARE study (NCT03017573), all treated with standard neoadjuvant chemotherapy (NAC) followed by surgery (Figure 1A and Table S1). For four patients, a third sample was collected at progression (P). Each case included detailed clinical annotation, enabling patient-resolved analysis of non-genetic ITH across therapy.

**Figure 1.**
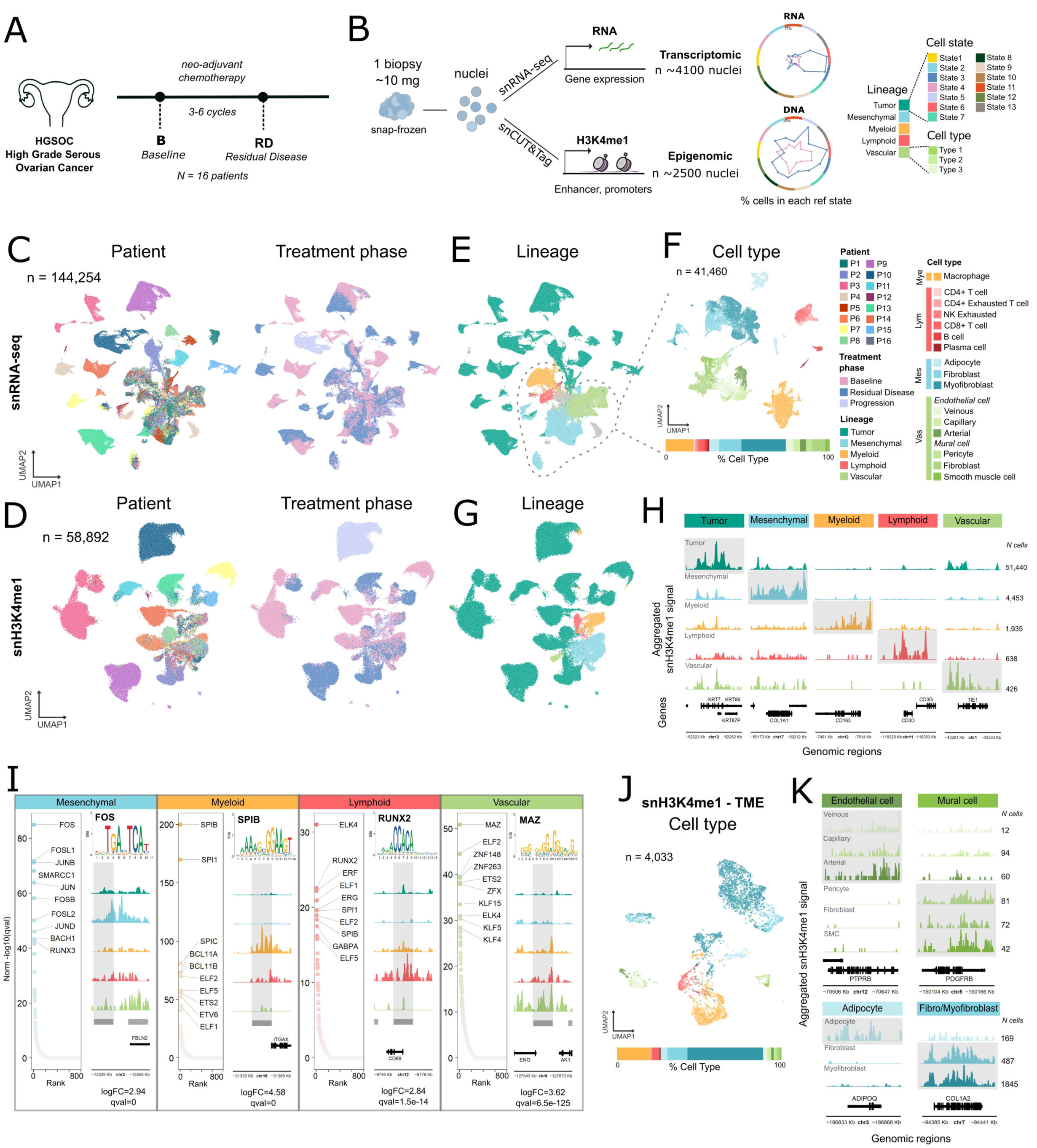
Single-nucleus multimodal profiling of ovarian cancer biopsies. (**A**) Overview of the SCANDARE clinical cohort, including treatment regimen and longitudinal biopsy collection at baseline (B) and residual disease (RD) following neoadjuvant chemotherapy (NAC). (**B**) Workflow illustrating single-nucleus RNA and histone (H3K4me1) profiling from milligram-scale frozen tumor biopsies and downstream computational analysis for cell type and state quantification. (**C** and **D**) UMAP projections of single-nucleus RNA (**C**) and H3K4me1 (**D**) datasets, with individual nuclei colored by patient identity or treatment phase (baseline or RD). (**E**) UMAP of snRNA-seq profiles colored by major cell lineages (epithelial, immune, stromal, endothelial). (**F**) Top: refined annotation of the tumor microenvironment (TME) showing distinct immune and stromal subtypes. Bottom: bar plot depicting their relative proportions across patients. (**G**) UMAP of snH3K4me1 profiles colored by lineage. (**H**) Aggregated snH3K4me1 signal tracks at representative marker genes illustrating lineage-specific enhancer activity. (**I**) Transcription factor (TF) motif enrichment in lineage-specific H3K4me1 peaks and representative genome tracks showing TF motifs within top-enriched regulatory regions. (**J**) Top: UMAP of TME nuclei from snH3K4me1 data, colored by cell type. Bottom: bar plot showing proportional abundance of each TME subtype. (**K**) Aggregated snH3K4me1 signal tracks at marker loci for vascular and mesenchymal cell types, highlighting enhancer-level regulatory diversity.

To allow parallel transcriptional and epigenomic analyses, we optimized nuclei extraction from snap-frozen, milligram-scale biopsies and established a workflow for single-nucleus RNA sequencing (snRNA-seq) and single-nucleus profiling of the histone modification lysine 4 mono-methylation at histone H3 (snH3K4me1) from the same material (Figures 1A and S1A). In contrast to snATAC-seq, which reveals chromatin accessibility, snH3K4me1 profiling shows enhancer- and promoter-associated histone marks that encode both active and poised regulatory regions. We optimized nuclei isolation and tagmentation conditions, benchmarking them against published protocols ^24,25^ for their ability to produce both high-quality snRNA and snH3K4me1 data (Figure S1B). Critical parameters included optimizing lysis buffer composition, nuclear dilution, and sample cleaning, aiming to preserve RNA integrity while allowing efficient *in situ* tagmentation of chromatin and maximizing nuclei yield (see the Materials and Methods).

We obtained high-quality snRNA-seq data for an average of 4,334 nuclei per biopsy (median 2,843 detected genes; 70% of nuclei with > 2,000 genes), which is comparable in coverage to fresh-tissue scRNA-seq data and the largest public HGSOC datasets (Figures 1C and S2A–S2B, and Table S2). For snH3K4me1 profiling, we recovered an average of 2,454 nuclei per biopsy (median 1,409 unique fragments per nucleus) (Figures 1D and S2C, and Table S3), generating one of the largest single-nucleus histone modification datasets from human tumors to date. These results demonstrate that integrated transcriptomic and epigenomic profiling is feasible from routine frozen biopsies, and that patient-resolved maps can be constructed for functional ITH across treatment.

### Single-nucleus profiling maps transcriptional and epigenomic diversity across tumor and microenvironmental compartments

snRNA-seq data from matched baseline and RD biopsies resolved the major lineages of the tumor and TME, including tumor epithelial, mesenchymal, myeloid, lymphoid, and vascular compartments (Figures 1E, 1F, and S2D-S2F). Within each lineage, refined cell types were readily distinguished using canonical markers. In addition to the expected TME populations (T cells, B cells, macrophages, fibroblasts), we also detected rare subtypes, such as plasma cells within lymphoid clusters (∼0.5% of TME cells per patient) and adipocytes within the mesenchymal compartment (∼4%), both of which were poorly resolved in scRNA-seq datasets. We also identified exhausted CD4⁺ T cells (CTLA4, TIGIT), exhausted NK cells (HAVCR2), and M2-polarized macrophages (SELENOP, MARCO). Importantly, snRNA-seq provided high-resolution delineation of vascular lineages, capturing arterial smooth muscle cells, pericytes, and adventitial fibroblasts, as well as arterial, capillary, and venous endothelial cells (Figures 1F, S2E, and S2F). This degree of vascular resolution is not achievable with standard scRNA-seq (Figure S2G-S2I), underscoring the utility of single-nucleus profiling for comprehensive TME characterization.

Complementary snH3K4me1 profiling captured enhancer- and promoter-associated histone landscapes across the same biopsies. Unlike snRNA-seq, which quantifies gene expression, snH3K4me1 measures enhancer and promoter-associated histone modifications and can be analyzed by gene- or peak-based partitioning (see Materials and Methods). Gene activity scores reflected lineage-defining marks, of tumor cells (KRT7, EPCAM), mesenchymal cells (COL1A1, DCN), and lymphoid cells (CD3D, CD2), allowing robust annotation of cell lineages (Figures 1G, 1H, S2J, and S2K). Differential peak analysis further identified lineage-specific enhancers, confirming activation of EMT and collagen pathways in mesenchymal cells and extensive enhancer priming of VEGF-related genes in vascular populations (Figure S2L). Motif enrichment analysis linked these enhancer signatures to lineage-defining transcription factors. SPI family factors were enriched in myeloid cells, while the MAZ motif was significantly enriched in regulatory regions of rare vascular cells (∼28 nuclei per patient; 426 total), demonstrating sensitivity sufficient to profile low-abundance populations (Figure 1I).

To systematically compare modalities, we transferred snRNA-seq annotations to snH3K4me1 data within the TME (Figures 1J, 1K, S2M, and S2N). All major and rare TME cell types were captured, including mural vascular cells and adipocytes. The strong alignment between transcriptional and chromatin-defined identities indicates that snH3K4me1 profiling resolves epigenomic heterogeneity with single-cell precision, thereby providing access to cell types not captured by bulk chromatin assays. Together, the transcriptomic and epigenomic datasets provide an integrated multimodal resource for dissecting non-genetic ITH across malignant and stromal compartments from milligram-scale biopsies.

### Consensus transcriptional programs define functional tumor states in HGSOC

We hypothesized, in line with previous studies ^26–28^, that tumor cells adopt a limited set of recurrent transcriptional programs—functional states—to adapt and survive in their local environment. While the events initiating these states may differ across patients, the resulting expression programs are often conserved. To identify functional recurrent tumor states in HGSOC, we assembled a discovery cohort by combining three HGSOC scRNA-seq datasets, two public ^29,30^ and one generated from fresh SCANDARE samples. This yielded a consensus atlas of 205,321 malignant cells from 65 patients and 150 samples (136 baseline and 14 RD; Table S4), which encompassed both primary and metastatic sites as well as tumors with homologous recombination deficiency (HRD) or proficiency (HRP) (Figures 2A and S3A).

**Figure 2.**
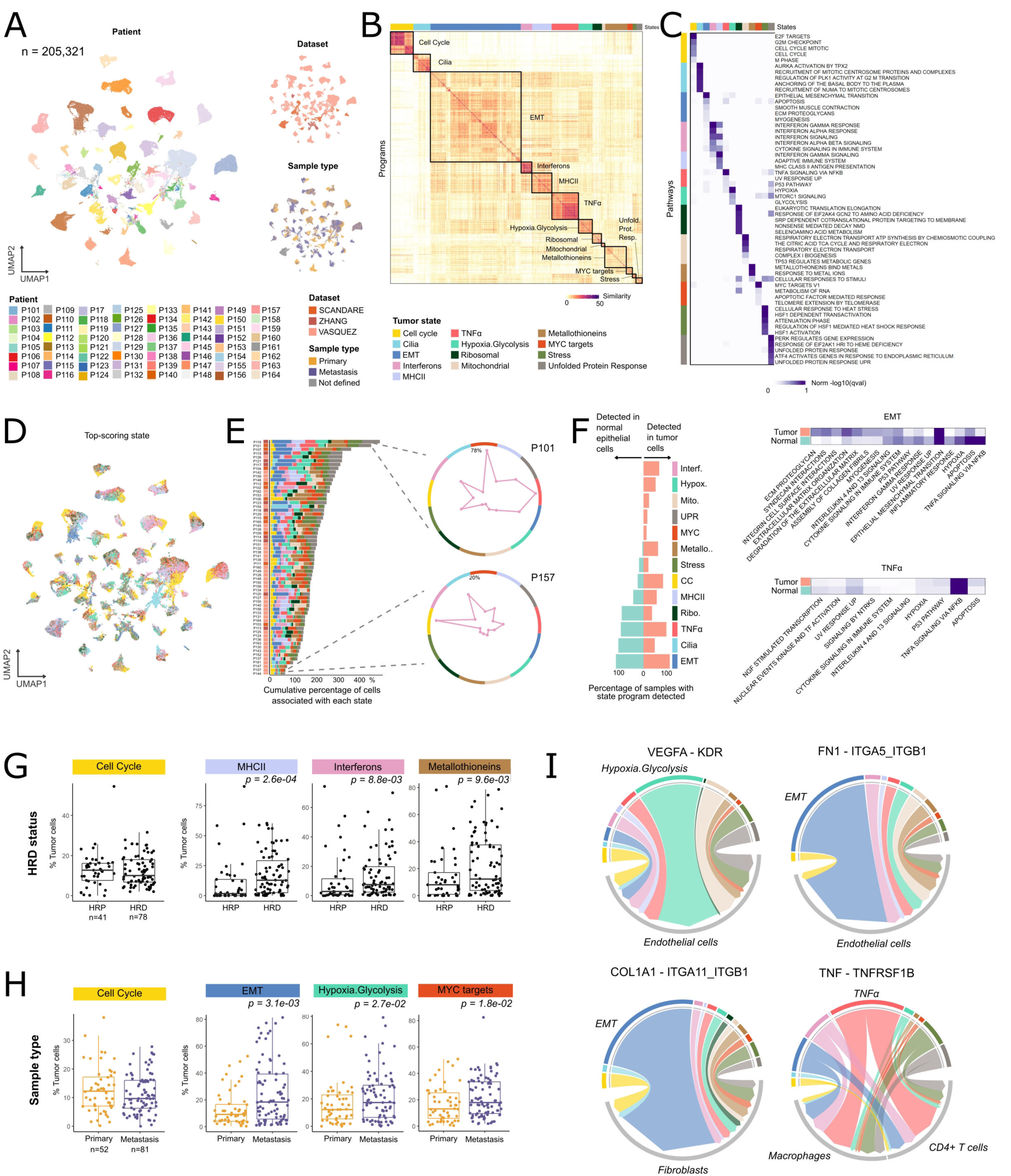
Consensus transcriptional programs define functional tumor states in HGSOC. (**A**) UMAP of tumor cells from integrated scRNA-seq datasets colored by patient identity, dataset origin, and sample type. (**B**) Pairwise overlap of top-scoring genes defining intratumoral expression programs across samples; hierarchical clustering reveals conserved transcriptional modules. (**C**) Pathway enrichment of the gene sets defining each module, used to annotate consensus tumor states. (**D**) UMAP of tumor cells colored by the dominant consensus state assigned through state-score decomposition. (**E**) Functional composition of baseline tumors: stacked bar plots show cumulative fractions of cells in each state per sample (left); representative radar plots illustrate state distributions for patients P101 (top) and P157 (bottom). (**F**) Comparison of normal and tumor epithelial samples: frequency of samples harboring each state (left) and pathway enrichment of EMT- and TNFα-associated programs derived separately from normal and tumor contexts (right). (**G** and **H**) Boxplots showing the fraction of tumor cells assigned to each state according to (**G**) homologous recombination deficiency (HRD) status or (**H**) anatomical site (primary vs metastatic). (**I**) Circos plots illustrating ligand–receptor interaction strength between tumor states and TME populations, highlighting state-specific signaling dependencies.

We applied non-negative matrix factorization (NMF) to decompose the transcriptional landscape of each tumor into shared gene expression programs (Figure 2B). Aggregating programs across patients defined thirteen robust consensus states, including proliferative, interferon, hypoxic, MYC, metallothionein, TNFα, and EMT-associated programs (Figures 2C and S3B, and Table S5). Several states aligned with metaprograms described in pan-cancer analyses, with EMT and cell cycle states encompassing multiple related modules (Figure S3C). We then functionally decomposed tumors by scoring individual cells for activation of each state, allowing cells to express multiple concurrent programs (Figures 2D and 2E). Despite genetic heterogeneity, these states were present in various combinations in all tumors, implying that they represent fundamental transcriptional modules conserved across HGSOC.

To contextualize these states, we compared them to expression programs in normal ovarian epithelium using the same method. Hypoxia and interferon-associated programs were tumor-specific, while EMT and TNFα-associated states were shared but transcriptionally rewired; for instance, cancer EMT programs were enriched for ECM remodeling genes (Figures 2F and S3D). We next explored associations between tumor cell states and available clinical features using the combined power of the three large cohorts (*n* = 136 baseline samples). As compared to HRP tumors, HRD tumors showed higher fractions of interferon-and MHC class II–associated states, consistent with increased inflammation in BRCA1-mutated cancers ^31^ and a higher mutational load and number of tumor-specific neoantigens in these tumors ^32^ (Figures 2G and S3E). Metastatic samples displayed higher representation of EMT, hypoxia, and MYC states as compared to primary sites (Figures 2H and S3F), suggesting that metastatic dissemination selects for transcriptional plasticity and angiogenic potential.

To assess the functional relevance of tumor states, we analyzed ligand–receptor interactions between malignant and microenvironmental cells (see Materials and Methods). Each state displayed a distinct signaling profile, indicating specialized modes of tumor–stroma communication (Figure 2I). EMT-, TNFα-, and hypoxia-associated cells showed the highest number of predicted interactions (Figure S3G). Hypoxic tumor cells preferentially signaled to endothelial cells through VEGFA/KDR, recapitulating canonical angiogenic crosstalk. EMT-enriched cells formed ECM–related contacts with fibroblasts (COL1A1/ITGA11) and endothelial cells (FN1/ITGA5), suggesting active remodeling of the TME. TNFα-high cells interacted with macrophages and CD4⁺ T cells via TNFR2/TNFRSF1B, a pathway known to promote immune suppression and drug resistance through recruitment of Tregs and tumor-associated macrophages ^33,34^. These findings link molecularly defined tumor states to distinct signaling dependencies within the HGSOC ecosystem, while validating our decomposition approach of determining ITH. Collectively, these analyses define a consensus map of HGSOC tumor states that integrates molecular and clinical diversity and provides a reference for quantifying functional ITH.

### Tracking tumor cell states under chemotherapy predicts chemoresistant evolution

Using our consensus reference, we functionally decomposed the tumor compartment for all SCANDARE samples profiled by snRNA-seq (Figure 3A). We scored each tumor cell for activation of the consensus states and compared cell state proportions between baseline and RD biopsies (Figures 3B, 3C, and S4A-S4C). States that decreased after treatment (logFC<0 and <10% of residual tumor cells) were defined as chemosensitive, while those that persisted or expanded were considered chemoresistant (logFC>0 and >10% residual cells). As expected, the cell cycle state was strongly depleted after therapy (median 15% in baseline, *vs* 1.5% in RD), consistent with targeting of proliferative cells by standard NAC regimens. In contrast, we observed increases in multiple patients for both TNFα signaling (with 6 of 16 patients showing >1-fold increase) and the EMT states (albeit with inter-patient variability) (Figure 3B).

**Figure 3.**
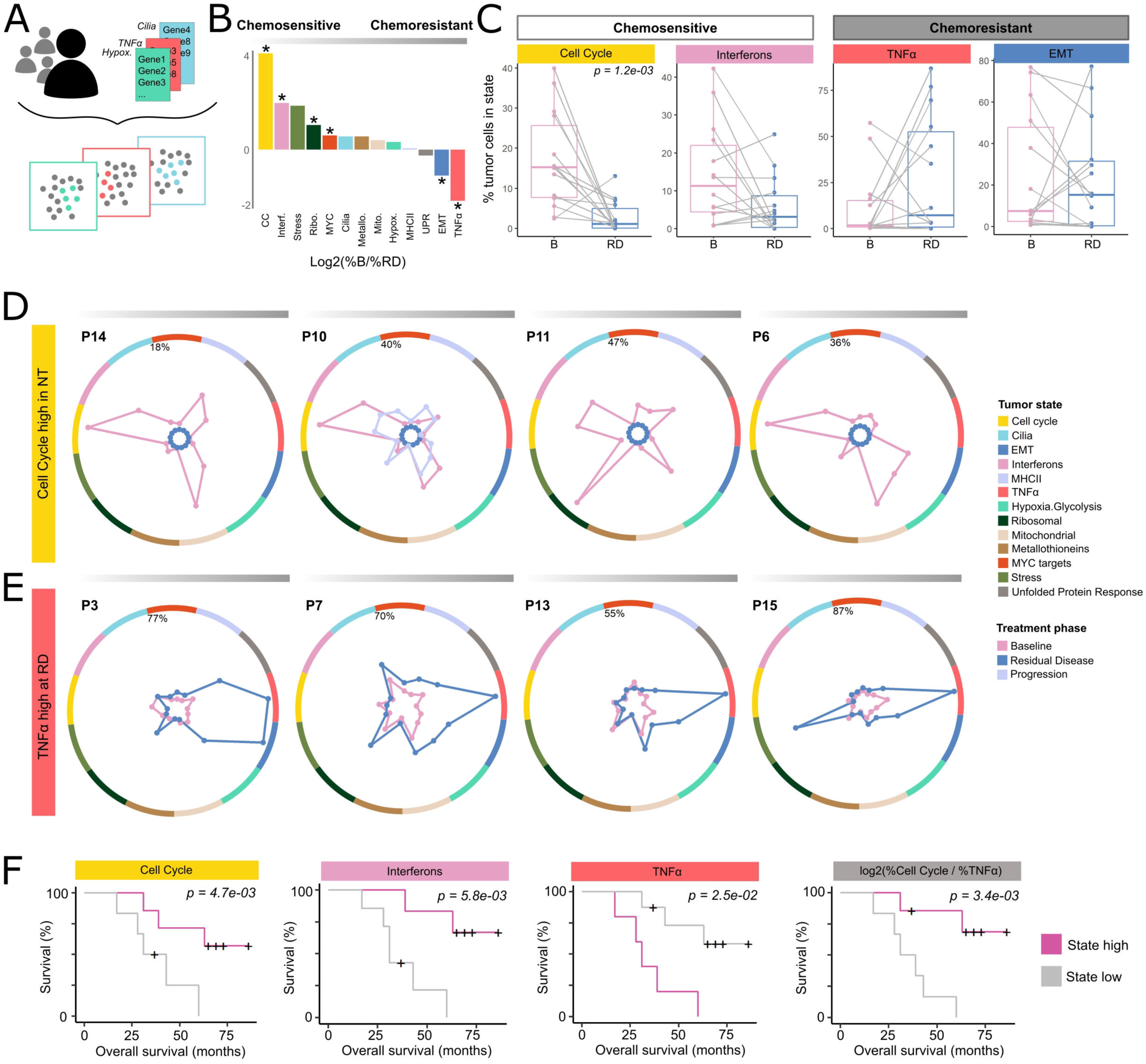
Tracking tumor cell states under chemotherapy reveals chemoresistant programs. (**A**) Schematic of the scoring framework for snRNA-seq data: each nucleus is independently scored for activation of consensus tumor-state signatures. (**B**) Log₂ fold-change in median fraction of cells per state between baseline (B) and residual-disease (RD) biopsies. Stars indicate chemosensitive (logFC<0 and <10% in RD) and chemoresistant states (logFC<0 and >10% in RD). (**C**) Paired boxplots comparing per-patient state frequencies before and after treatment. (**D** and **E**) Representative radar plots showing functional tumor composition for individual patients, where each axis corresponds to a consensus tumor state ordered from chemosensitive (left) to chemoresistant (right). Patients in (**D**) exhibit high baseline cell cycle activity and strong treatment response, whereas patients in (**E**) show post-treatment expansion of TNFα-associated cells in RD samples. (**F**) Kaplan–Meier survival curves stratified by baseline abundance of tumor cells in the cell cycle, interferon, or TNFα states, and by the log-ratio of cell cycle to TNFα state fractions, linking baseline functional composition to clinical outcome.

A paired statistical analysis (N = 14 patients) confirmed a significant treatment-associated depletion of the cell cycle program (*p* < 0.01). To strengthen these findings, we integrated the paired samples present in our reference HGSOC scRNA-seq atlas (N = 11 additional matched pairs from ^30^ and the fresh SCANDARE samples, see Figure 2) with frozen samples. This analysis reinforced our findings, showing significant decreases in individual patients post-treatment in the four chemosensitive cell states: cell cycle, MYC, ribosomal and interferons associated programs (Figure S4A). Notably, MYC-associated programs declined post-treatment, in line with the role of MYC in proliferation. The hypoxia-associated state was partially responsive to treatment, suggesting reduced pro-angiogenic activity. In contrast, TNFα activation strongly increased after NAC (median 34% in RD *vs* 7.1% in baseline), and the EMT state either remained unchanged or increased, supporting a model in which inflammatory and mesenchymal programs mark chemoresistant subpopulations. Note that the stress-associated program was detected far less using snRNA-seq on frozen biopsies (2.7% median) than with scRNA-seq on fresh specimens (13.7% median). However, as this pattern is consistent with well-known dissociation-induced stress artifacts in fresh tissue, it likely reflects a technical effect rather than a true biological difference.

Longitudinal changes in individual patients revealed recurrent patterns (Figures 3D, 3E, and S4D). Radar plots visualizing the fraction of cells activating each state showed that responders exhibited flat RD profiles with no detectable tumor states, while their baseline tumors were enriched in cell cycle and interferon programs (Figure 3D). In contrast, partial responders frequently had RD biopsies enriched for the TNFα/EMT resistant states (Figure 3E). Some partial responders already exhibited high TNFα/EMT states at baseline, indicating pre-existing resistance states (Figure S4D). In one patient with serial sampling (patient P10; Figure 3D), the progression tumor resembled the baseline composition but contained fewer cycling cells (40% vs. 13%; *p* = 2 × 10⁻¹⁵⁰), suggesting relapse through re-emergence of a less proliferative, therapy-tolerant clone.

Critically, baseline compositions correlated with outcome: tumors enriched for cell cycle or interferon-responsive states were associated with longer overall survival, and those enriched for TNFα-positive cells, with a poor prognosis (Figure 3F). A simple baseline ratio of cell cycle/TNFα states improved survival discrimination as compared to taking each percentage separately (*p* = 0.0043). These results demonstrate that our ITH scoring framework, based on single-nucleus profiling, can monitor functional ITH from frozen biopsies and identify therapy-resistant tumor subpopulations linked to clinical response.

### Epigenomic landscapes encode chemoresistant states prior to therapy

We next used snH3K4me1 profiling to examine how these functional states are epigenetically encoded. We computed the *epigenomic cell state* scores for each nucleus to quantify how strongly H3K4me1 marked the enhancer regions of consensus state genes in individual cells (Figure 4A). As in transcriptomic data, cells frequently had concurrent priming for multiple states of epigenomic patterns (Figures 4B and S5A). Across patients, subsets of tumor cells displayed strong epigenomic priming at state-specific loci (Figure 4C), indicating that the histone modification landscapes partially encode functional states.

**Figure 4.**
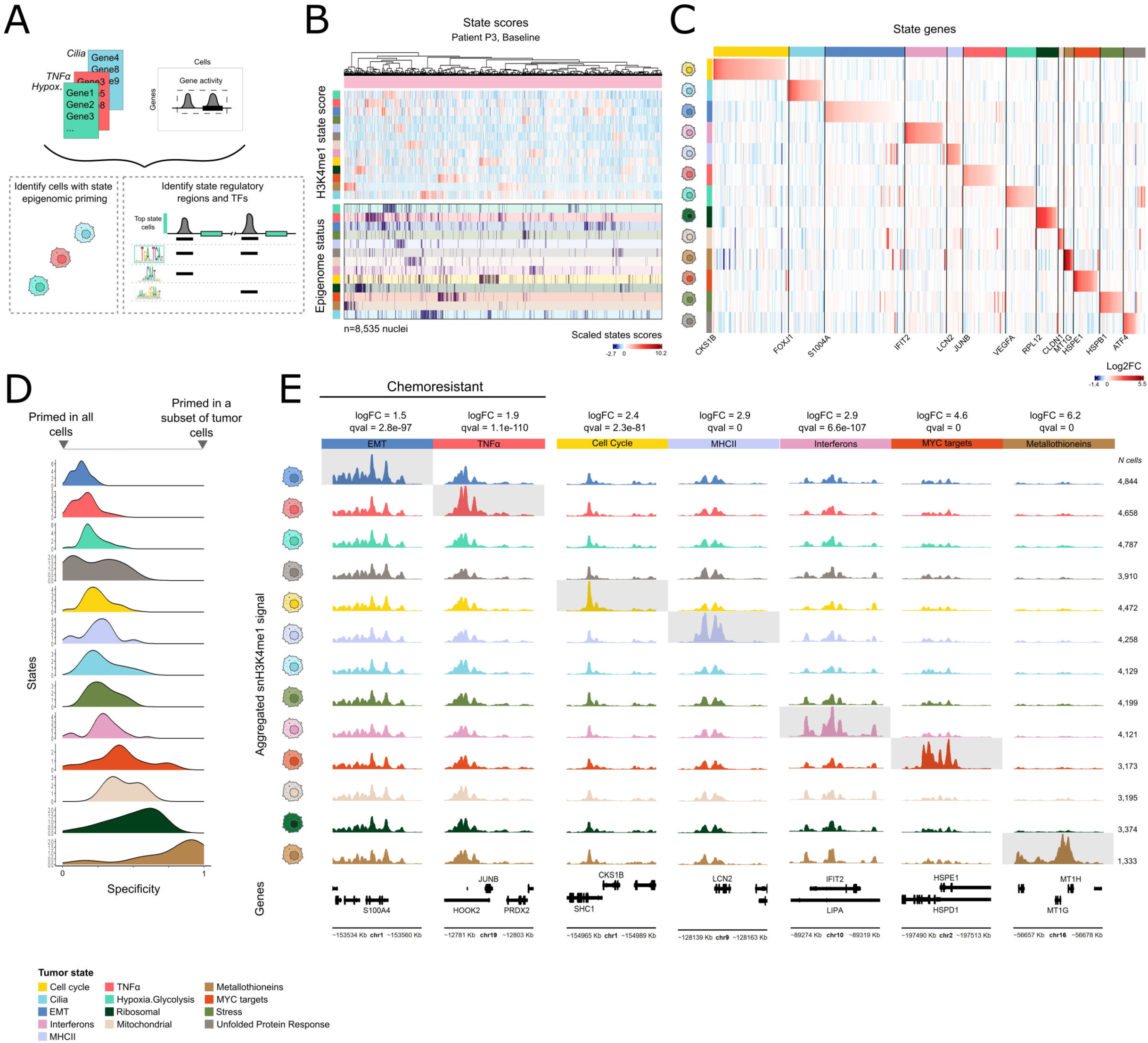
Enhancer landscapes encode functional tumor states in HGSOC. (**A**) Schematic of the computational workflow used to derive epigenomic state scores from single-nucleus H3K4me1 profiles. (**B**) Heatmaps for patient P3 at baseline showing per-cell epigenomic state scores (top) and resulting state assignments (bottom). (**C**) Log₂ fold-change in H3K4me1 signal for each state compared with all others, displayed across genes defining the consensus tumor-state programs. (**D**) Distribution of gene-level specificity scores illustrating the extent of enhancer priming for each state. (**E**) Representative genome tracks showing aggregated snH3K4me1 signal across nuclei from individual states, with corresponding log₂ fold-change and q values for differential activity indicated above each locus.

To assess the specificity of this priming, we calculated an *epigenomic specificity score* for each gene state (Figure 4D). Genes with high specificity, such as MT1G and MT1H, were enriched in a restricted subset of cells, whereas broadly primed (low specificity) genes, such as canonical EMT or TNFα components (e.g., S100A4 and JUNB), showed H3K4me1 marking in a large fraction of tumor cells (Figures 4E and S5B). Most states (9 of 13) exhibited specific priming (> 1.5-fold enrichment for > 50% of their genes) (Figure S5C). Genes defining the MYC-associated program were highly cell-specific (with 80% of genes specifically primed). In stark contrast, the EMT and TNFα programs were broadly H3K4me1-primed (with specific priming in only 17% of TNFα genes, and 7% of EMT genes), implying that these programs are epigenetically poised throughout the tumor and consistent with their role as resistant states. Such genome-wide priming may enable rapid activation under stress, akin to bivalent chromatin states in pluripotent cells, where H3K27me3-repressed loci remain pre-marked with H3K4me1/3 ^35^.

### Transcription factor networks organize enhancer-primed resistant states

As the TNFα and EMT programs were broadly primed in all tumor cells yet only over-expressed in a subset, we hypothesized that chemotherapy selects or activates cells through specific transcription factor (TF) networks. To identify candidate regulators, we analyzed H3K4me1 peaks (mean 1.5 kb, ∼75 per tumor state on average) associated with state-specific genes. Motif enrichment revealed recurrent TF families enriched at regulatory regions for each cell state. We identified several known regulators of tumor functions, including HIF1A for the hypoxia-associated cell state and members of the IRF TF family (IRF2, 7, 8 and 9) for the interferon cell state (Figures 5A and S6A).

**Figure 5.**
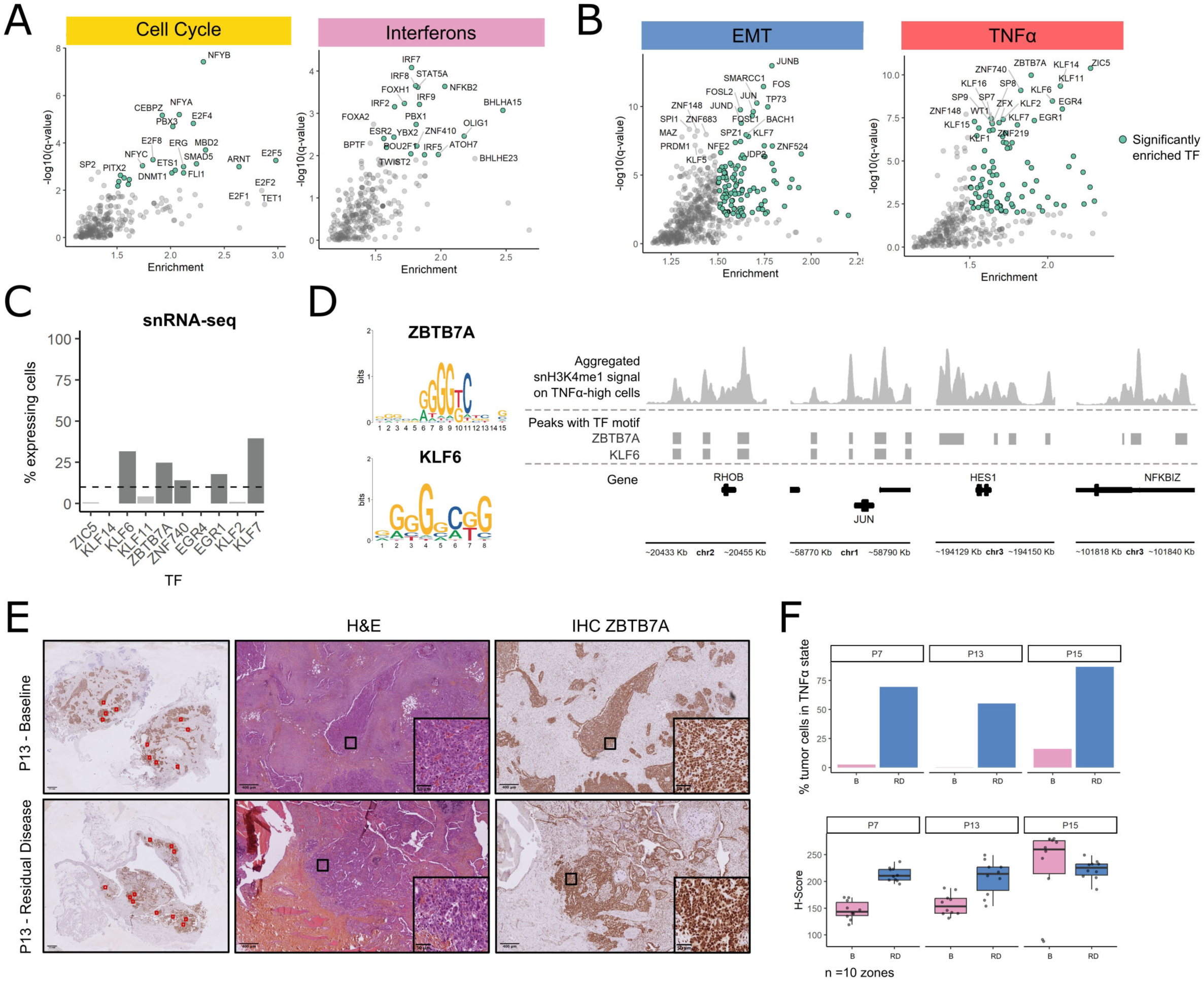
Transcription-factor networks organize enhancer-primed chemoresistant states. (**A** and **B**) Scatter plot of predicted transcription-factor (TF) binding near tumor state genes, showing –log₁₀(q value) vs. enrichment fold-change from snH3K4me1 data, for (**A**) chemosensitive states (cell cycle and interferons) and (**B**) chemoresistant states (EMT and TNFα). (**C**) Fraction of tumor cells expressing the top 10 predicted TFs according to snRNA-seq. (**D**) Genome tracks showing aggregated snH3K4me1 profiles in TNFα-assigned cells at regulatory peaks containing ZBTB7A and/or KLF6 motifs. (**E**) Immunohistochemistry (IHC) for ZBTB7A on paired FFPE tumor sections from patient P13 collected at baseline and after chemotherapy. (**F**) Quantification of ZBTB7A staining intensity (H-score) across ten independent tumor regions for patients P7, P13, and P15, demonstrating increased protein abundance in post-treatment samples.

Candidate drivers were defined as factors with (i) motif enrichment near state genes and (ii) specific expression within the corresponding tumor cells. For the chemoresistant TNFα-associated state, the top TF candidates included KLF6, ZBTB7A, ZNF740, EGR1, and KLF7 (Figures 5B and 5C). The ZBTB7A motif occurred near 11 of 39 TNFα-program genes and was expressed in 25% of tumor cells, while KLF6 and EGR1 were expressed in 32% and 18% of tumor cells (Figure 5C), respectively, with motif enrichment in 9 of 39 TNFα-program genes. ZBTB7A was predicted to regulate multiple key genes of the TNFα inflammation pathway, including ATF3 and NFKBIZ in the NF-κB signaling cascade (Figure 5D). Notably, analysis of ZBTB7A via immunohistochemistry on paired FFPE tumor sections adjacent to frozen samples confirmed its increased expression in residual tumors from 2 of 3 patients who exhibited TNFα-state expansion (Figures 5E, 5F, S6B, and S6C); in the third case, uniformly high baseline levels were maintained. Thus, ZBTB7A expression increased following treatment in parallel to an expansion of cells in a TNFα state.

### Chemotherapy remodels fibroblast enhancer programs

We next examined microenvironmental evolution during chemotherapy. Transcriptomic analysis showed major shifts in the TME composition (Figures 6A, 6B, and S7A). Across paired samples, fibroblasts increased from 2% to 11% of the TME cells (*p* = 0.011), accompanied by higher infiltration of CD4⁺ and CD8⁺ T cells (2.4- and 1.7-fold increases, respectively) (Figures 6B and S7B). These findings parallel reports of increased lymphocytic infiltration in residual triple-negative breast cancers associated with poor outcome ^36^. Conversely, vascular capillary and pericyte populations declined (*p* = 0.024 and 0.016, respectively), suggesting decreased pro-angiogenic activity post-therapy. Such findings echo intrinsic tumor evolution patterns found above, with a decrease upon treatment in tumor cells in pro-angiogenic state. Patients with HGSOC are often treated with an anti-angiogenic drug (bevacizumab) in an adjuvant setting; however, our data suggest that it might be more effective prior to NAC, as baseline pre-NAC tumors appear to be more dependent on VEGFA signaling than residual tumors. Further, we found that baseline biopsies with higher fibroblast content correlated with worse overall survival, and those with CD4⁺ and NK-cell enrichment, with favorable outcomes (Figure S7C).

**Figure 6.**
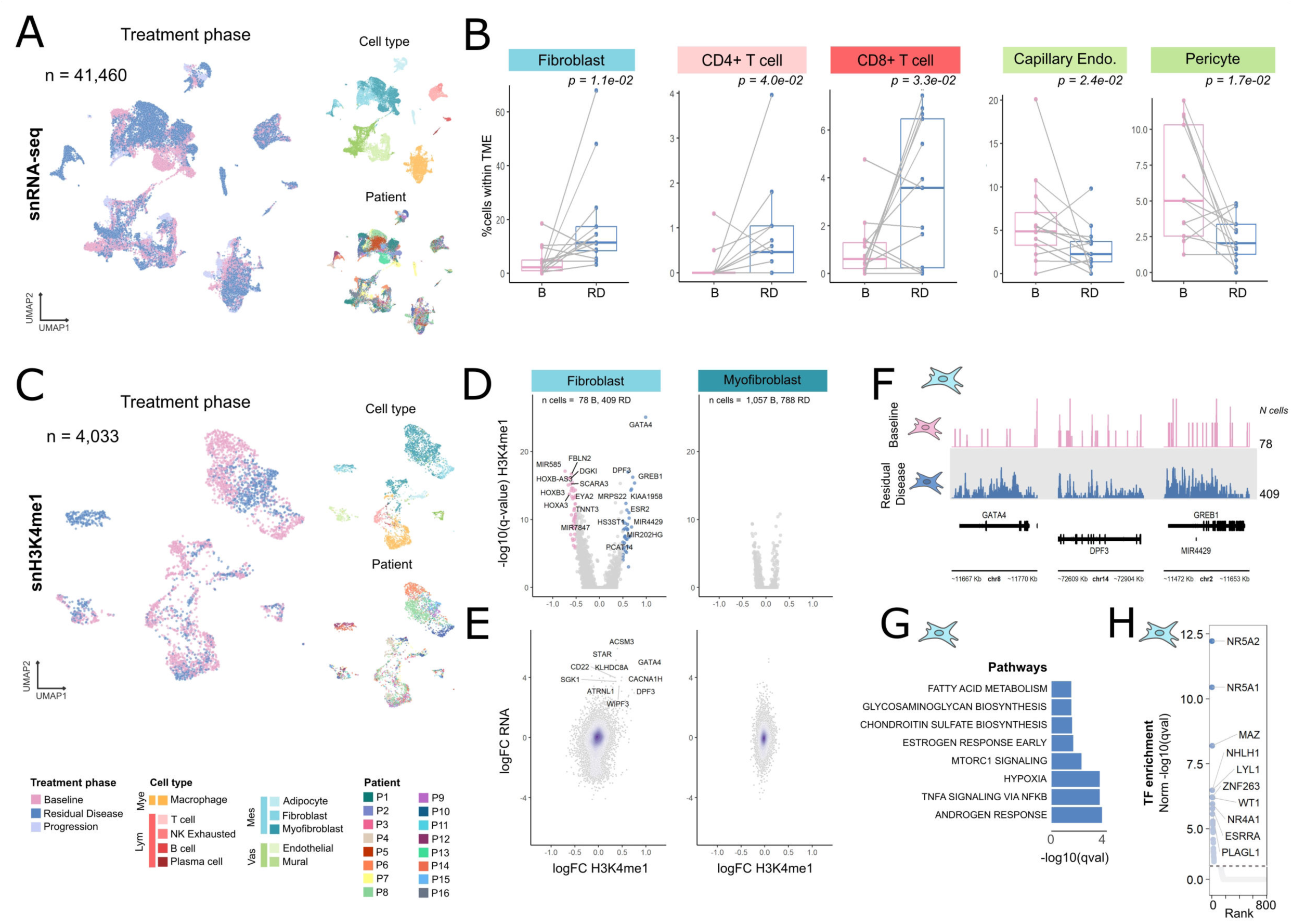
Chemotherapy remodels the tumor microenvironment and fibroblast enhancer programs. (**A**) UMAP of TME nuclei from snRNA-seq data colored by treatment status, cell type, and patient. (**B**) Paired boxplots showing the proportion of each TME cell type before (baseline, B) and after treatment (residual disease, RD) across patients. (**C**) UMAP of snH3K4me1 data colored by treatment phase, cell type, and patient. (**D**) Volcano plot of differential H3K4me1 activity across cell types between baseline and RD samples (Wilcoxon test). (**E**) Concordance of transcriptional and epigenomic remodeling: density plot of log₂ fold-changes in snRNA-seq (y-axis) vs. snH3K4me1 (x-axis) for matched cell types. (**F**) Genome tracks displaying aggregated H3K4me1 signal in fibroblasts from baseline and RD samples at loci with enhancer gain in RD fibroblasts. (**G**) Pathway enrichment for genes up-regulated in fibroblasts after chemotherapy based on pseudo-bulk snRNA-seq analysis. (**H**) Scatter plot showing TF-motif enrichment in top-specific peaks from RD fibroblasts, highlighting candidate regulators of fibroblast reprogramming.

Finally, paired transcriptomic–epigenomic comparisons revealed stromal remodeling dominated by fibroblasts (Figures 6C–6F and S7D–S7F). Upon treatment, > 84 loci showed H3K4me1 remodeling specifically in fibroblasts, whereas macrophages and myofibroblasts remained stable (Figures 6D and S7F). Genes gaining H3K4me1 signal were transcriptionally upregulated in RD tumors (Figure 6E), implicating activation of TNFα and hormone sensing pathways (androgen and estrogen; Figure 6G). Motif analysis of remodeled enhancers identified NR5A1 and NR2F2 as potential regulators of these transitions (Figure 6H). Thus, chemotherapy reshapes the TME at both the cellular and epigenomic levels, with pronounced remodeling of the fibroblast compartment. Tumor and stromal cells undergo parallel epigenomic evolution, together establishing a chemoresistant ecosystem.

## Discussion

Our work provides proof of concept that functional ITH can be monitored at high resolution from frozen biopsies collected during routine clinical care. Whereas previous single-cell histone profiling studies relied on model systems or large amounts of input material ^22,37–39^; our multimodal single-nucleus approach enables *in situ* tagmentation of histone modifications from low-input, clinically-relevant specimens. We find that H3K4me1 marks stratify both TME and malignant subpopulations, and that specific regulatory elements are linked to chemoresistance. By combining single-nucleus RNA and histone profiling, we captured both the transcriptional and the epigenomic dimensions of tumor evolution in patients with HGSOC. Our results highlight that epigenetic diversity is a critical indicator of tumor ecosystems. Notably, we found that chemoresistant tumor cell states do not simply expand post-treatment but rather already exist in an epigenetically primed configuration prior to treatment in cancer cells. Thus, pre-existing chromatin states could contribute to tumor adaptability and relapse and could represent targets for adjuvant strategies.

In model systems, chromatin variation generates metastable cell states that enable stress tolerance and phenotypic plasticity ^7,8,10^. Here, we demonstrate this principle in human tumors through longitudinal sampling. Chemoresistant states marked by TNFα signaling and EMT are epigenetically primed through H3K4me1 modifications prior to therapy but are transcriptionally active only in subsets of malignant cells. These programs align with inflammation-driven mesenchymal pathways linked to resistance across cancers ^40,41^. Their poised landscapes likely facilitate rapid adaptation to therapeutic stress, explaining the persistence of resistant subpopulations. Indeed, chemotherapy-induced stress triggers transcriptional activation in triple-negative breast cancer models exposed to cytotoxic agents^22^.

We established a unified approach to quantify non-genetic ITH in HGSOC by defining the functional states that tumor cells adopt within the TME. Many states overlapped with those observed in other epithelial cancers ^26,28^, and EMT emerged as a recurrent signature, integrating multiple EMT-related metaprograms. Recent studies indicate that mitochondrial-high malignant cells may reflect metabolic dysregulation relevant to chemoresistance ^42^. Thus, while previous metaprograms have excluded cell states with high mitochondrial gene expression as technical artifacts, we included their quantification and propose that they be evaluated as potential metabolic biomarkers of treatment outcome. Our integrative chromatin–transcriptome profiling revealed the TF ZBTB7A as a likely regulator of the TNFα-associated chemoresistant states, and these findings are supported by increased ZBTB7A protein levels in residual tumors. These results suggest that chromatin–transcriptome profiling can directly identify candidate regulators from patient material.

Our patient-resolved ITH maps capture not only the malignant plasticity but also the microenvironmental co-evolution, whereby loss of interferon and hypoxia signatures parallels expansion of inflammatory and mesenchymal states. Notably, tumor compositions at baseline strongly correlated with the patient outcome after treatment. Tumors enriched in proliferative and interferon-responsive states were associated with an improved prognosis, whereas those dominated by TNFα programs were associated with poor survival, consistent with previous studies ^43^. The TNFα–TNFR2/TNFRSF1B axis emerged as a major signaling route of chemoresistance ^33,34^, in line with evidence that TNFR2 inhibition can restore immune activation and enhance tumor clearance ^44^. Conversely, interferon signaling and cell cycle activity appeared protective, suggesting that maintaining proliferative competence or enhancing interferon responses could improve response to therapy.

Chemotherapy also remodeled the TME fibroblast population, which underwent recurrent expansions and extensive epigenomic reprogramming via TNFα and hormone-related pathways. This stromal adaptation likely supports chemoresistant tumor states through reciprocal signaling, emphasizing that therapy resistance arises from coordinated tumor–stroma evolution rather than cell-autonomous mechanisms alone. Overall, tracking TME transitions in serial biopsies provides a quantitative view of tumor evolution under therapy. Profiling baseline tumors for transcriptomic and epigenomic composition could identify patients predisposed to relapse and provide a guide for an earlier intervention with intensified or epigenetic therapies to limit plasticity ^45^. Systematic tumor cell state decomposition may therefore serve both as a biomarker and as a framework for predicting adaptive trajectories.

This work establishes a generalizable strategy for integrating multimodal single-nucleus profiling into oncology and for tracking tumor evolution in clinical cohorts. Because the workflow requires only milligram-scale frozen biopsies, it is readily transferable to other solid tumors and suitable for longitudinal and retrospective studies. The resulting resource—including the consensus reference atlas, patient-level datasets, and analysis code—provides a practical foundation for dissecting chromatin-encoded plasticity in human cancers and for designing precision approaches informed by functional ITH.

---

**Figure S1. Optimization of nuclei extraction and multimodal library generation.** (**A**) Detailed schematic of the workflow for processing milligram-scale snap-frozen tumor biopsies for single-nucleus RNA and H3K4me1 profiling. (**B**) Comparison of alternative nuclei-extraction protocols showing nuclei yield, snRNA-seq library complexity, and snH3K4me1 snCUT&Tag library quality, highlighting the optimized method used in this study.

**Figure S2. Quality control and annotation of multimodal single-nucleus datasets.** (**A** to **C**) Quality-control metrics for snRNA-seq, scRNA-seq, and snH3K4me1 samples showing numbers of nuclei, detected genes or fragments, and mitochondrial or peak percentages. (**D** to **G**) Cell lineage and cell type distributions annotated by canonical markers. (**H** to **I**) Validation across SCANDARE scRNA-seq and reference datasets. (**J** to **L**) Lineage-resolved snH3K4me1 enhancer activity and pathway enrichments. (**M**) Joint UMAP embedding by CCA alignment of snRNA-seq and snH3K4me1 data. (**N**) Cell type composition of snH3K4me1 data across samples.

**Figure S3. Definition and validation of consensus tumor states.** (**A**) Overview of the integrated scRNA-seq cohort and processing pipeline for state discovery. (**B** to **D**) Pathway enrichments, gene overlaps, and state gene intersections for EMT and TNFα programs. (**E** and **F**) Boxplots of state proportions by HRD status and anatomical sampling site. (**G**) Bar plots quantifying numbers of ligand–receptor interactions per tumor state as sender (top) or receiver (bottom).

**Figure S4. Chemotherapy-associated remodeling of tumor state composition.** (**A** to **C**) Paired boxplots showing per-state frequencies between baseline and residual samples from paired snRNA-seq and scRNA-seq datasets. (**D**) Radar plots of per-patient tumor compositions ordered from the most chemosensitive (left) to the most chemoresistant (right) states.

**Figure S5. Epigenomic encoding of tumor state identity.** (**A**) Epigenomic state activity matrices and cell state assignments for representative baseline (B) samples (from patients P7, P8, and P14). (**B**) Aggregated snH3K4me1 signal tracks around state-defining genes, with log₂ fold-change and q values indicated. (**C**) Fraction of state genes with log₂ fold-change > 1.5, indicating state-specific enhancer priming.

**Figure S6. Identification of transcription-factor regulators of TNFα-associated states.** (**A**) Volcano plots of TF-motif enrichment near genes of each tumor state from snH3K4me1 data. (**B** and **C**) Immunohistochemistry for ZBTB7A in FFPE tumor sections from patients P7 and P15 before and after chemotherapy.

**Figure S7. Chemotherapy-induced remodeling of the tumor microenvironment.** (**A** and **B**) TME cell counts and proportions by treatment phase from snRNA-seq data. (**C**) Kaplan–Meier survival curves for patients stratified by baseline fibroblast, CD4⁺ T-cell, and exhausted NK-cell abundance. (**D** to **F**) Differential expression and enhancer activity analyses of TME cell types between baseline and residual samples across modalities.

